# A comprehensive benchmark of publicly available image foundation models for their usability to predict gene expression from whole slide images

**DOI:** 10.64898/2026.03.02.709012

**Authors:** Arfa Jabin, Shandar Ahmad

## Abstract

Recent advances in large-scale self-supervised learning have led to the emergence of foundation models capable of extracting transferable visual representations from high-dimensional image data. In computational pathology, such models are increasingly used as feature encoders for molecular prediction tasks. However, systematic benchmarking of publicly available image foundation models for transcriptomic prediction from whole-slide images (WSIs) remains limited. Here, we perform a comprehensive evaluation of five state-of-the-art vision foundation models-DINOv2, Phikon, UNI, H-Optimus-0, and MedSigLIP-for gene expression prediction using the TCGA-BRCA cohort. Tile embeddings extracted from each model were aggregated via attention-based multiple instance learning (MIL), followed by multi-target regression to predict RNA-seq expression profiles. Performance was assessed using gene-level Spearman correlation across samples. Histopathology-specific foundation models consistently outperformed general-purpose encoders, with Phikon achieving the strongest overall performance, followed by UNI and H-Optimus-0. These findings demonstrate that domain-aligned pretraining substantially enhances morphology-to-transcriptome inference and provide a principled benchmark for foundation model selection in molecular pathology.

## 1. Introduction

The digitization of histopathological slides into gigapixel-scale whole-slide images (WSIs) has transformed diagnostic pathology into a computational discipline capable of large-scale quantitative analysis. Deep learning approaches now enable extraction of morphological representations that correlate with molecular phenotypes, including gene expression and pathway activity (Coudray et al., 2018; He et al., 2020; Schmauch et al., 2020). Parallel to this shift, large-scale self-supervised foundation models have redefined visual representation learning (Krizhevsky et al., 2012; He et al., 2016; Dosovitskiy et al., 2021; He et al., 2022). These models learn general-purpose embeddings through contrastive learning, masked modeling, or self-distillation objectives (Devlin et al., 2019; Radford et al., 2021). In digital pathology, domain-specific foundation models trained on millions of histology patches have emerged as powerful feature extractors capable of capturing hierarchical morphological structure.

Despite widespread adoption, systematic benchmarking remains tentatoive, particularly when downstream tasks differ from pretraining objectives (Raghu et al., 2019; Kornblith et al., 2019). Gene expression prediction from WSIs represents a stringent test of representation quality, requiring sensitivity to subtle morphology-linked transcriptomic variation.

Here, we benchmark publicly available foundation models spanning general vision and pathology-specific pretraining regimes to quantify their suitability for transcriptomic inference from histology.

## 2. Materials and Methods

### 2.1 Dataset and Cohort Description

#### 2.1.1 Data sources and inclusion criteria

We utilized the TCGA Breast Invasive Carcinoma (TCGA-BRCA) cohort generated by The Cancer Genome Atlas Network (2012) and further described within the Pan-Cancer framework by Weinstein et al. (2013). Data were accessed through the NCI Genomic Data Commons (GDC), which provides harmonized molecular and imaging datasets (Grossman et al., 2016). The TCGA-BRCA cohort includes matched Hematoxylin and Eosin (H&E)-stained diagnostic whole-slide images (WSIs) and bulk RNA sequencing (RNA-seq) profiles for primary breast tumors. The cohort captures diverse molecular subtypes of breast cancer, including hormone receptor–positive, HER2-enriched, and triple-negative tumors, reflecting established intrinsic subtype classifications (Perou et al., 2000; Parker et al., 2009). This diversity enables investigation of transcriptomic–morphologic associations across biologically heterogeneous disease states.

#### 2.1.2 Cohort Assembly

RNA-seq data were available for 1,231 primary tumor cases. Intersection with available diagnostic histology resulted in 1,193 candidate cases with matched WSI and RNA-seq data. After quality control, file verification, and exclusion of slides with insufficient tumor tissue or processing artifacts, 987 cases remained with complete, usable WSI and RNA-seq profiles for downstream analysis. Whole-slide images were provided in SVS format at 40× magnification. To maintain a one-to-one correspondence between morphology and transcriptome, a single representative diagnostic slide was selected per patient when multiple slides were available.

### 2.2 RNA-seq Processing and Normalization

RNA-seq reads were aligned using the STAR aligner (Dobin et al., 2013) under the GDC harmonized workflow. Gene-level expression quantification was derived using FPKM-UQ (upper-quartile normalized fragments per kilobase per million mapped reads), consistent with TCGA processing conventions (Grossman et al., 2016). Overall, RNA-seq expression profiles comprised approximately 60,000 genes per sample. Expression matrices were filtered to retain valid Ensembl gene identifiers and to remove technical artifacts. Log transformation was applied for variance stabilization. Subsequently, min–max normalization was performed to scale gene expression values and improve numerical stability across genes. The resulting normalized expression values served as slide-level supervision signals for downstream modeling.

### 2.3 Foundation Models Evaluated

We benchmarked five publicly available encoders representing distinct pretraining paradigms:

- **DINOv2** (Oquab et al., 2023) - self-supervised ViT pretrained on natural images.
- **Phikon** (Filiot et al., 2024) - pathology-specific DINO-based model trained on pan-cancer histology.
- **UNI** (Chen et al., 2024) - large-scale pathology model trained on >100M histology patches.
- **H-Optimus-0** (Karasikov et al., 2025) - billion-parameter ViT-g pathology model.
- **MedSigLIP** (Zhai et al., 2023) - medical vision-language pretraining framework.

These models span general vision, multimodal, and domain-specialized histopathology pretraining regimes (see Table 1).

**Table 1.**
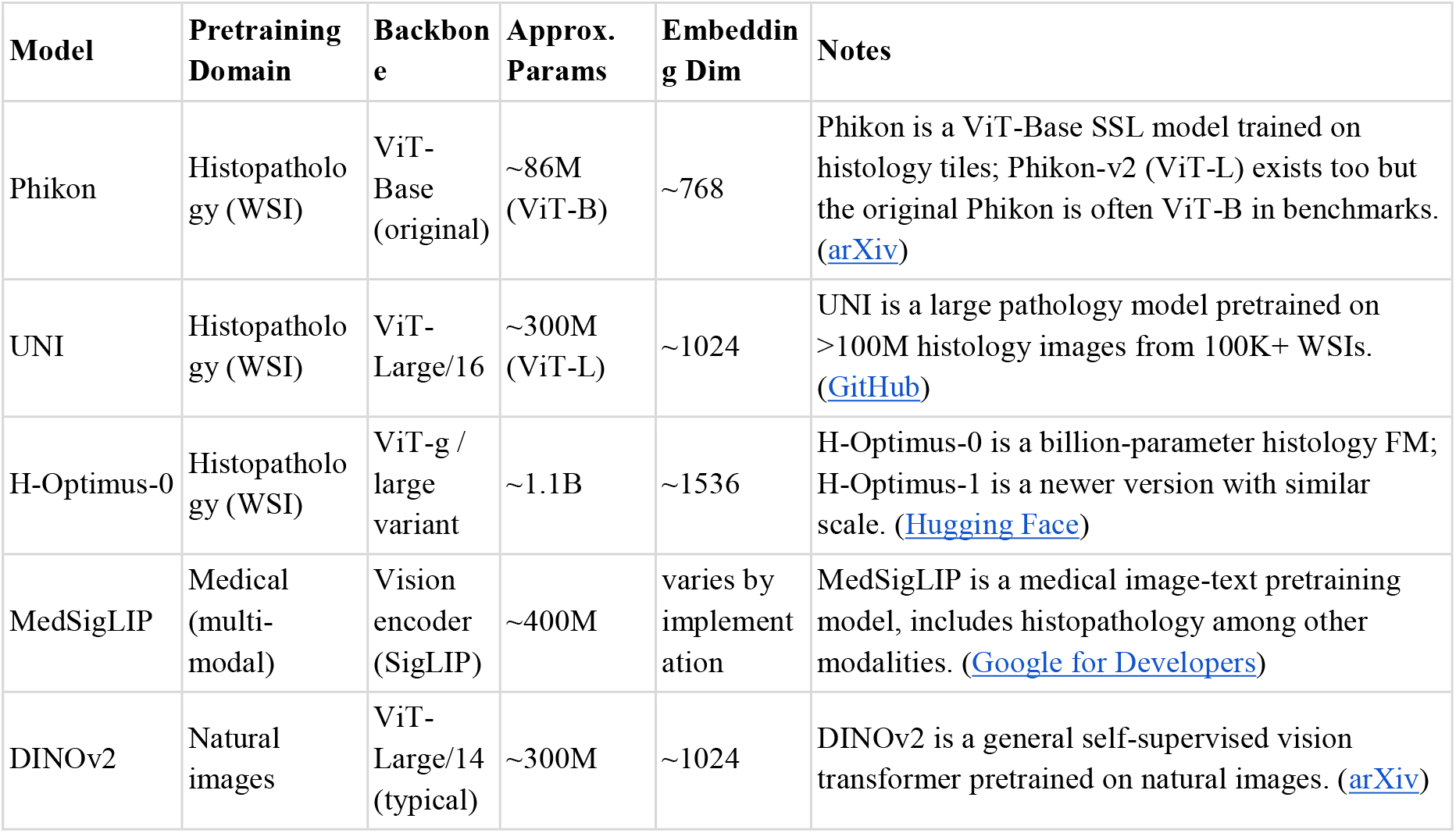
Foundation Models characteristics.

### 2.4 Prediction Framework

WSIs were partitioned into tiles. For each model, frozen tile embeddings were extracted (e.g., 768–1536 dimensional vectors depending on encoder). An attention-based multiple instance learning (MIL) framework aggregated tile embeddings into slide-level representations. The aggregated embedding was passed through a fully connected regression head to predict continuous gene expression values via multi-target regression. (Ilse, Tomczak, & Welling, 2018). Model optimization minimized prediction error between predicted and observed RNA-seq values.

### 2.5 Evaluation Metrics

Performance was quantified using gene-level Spearman correlation (ρ) across samples. Additional analyses included:

- Distributional comparison of gene-level ρ
- Empirical cumulative distribution functions (ECDF)
- Rank-based correlation curves
- Threshold-based summaries (ρ > 0.3 and ρ > 0.5)

These complementary analyses ensured robust cross-model comparison.

## 3. Results

### 3.1 Foundation Model Benchmarking

We evaluated gene expression prediction performance across five WSI feature extractors: DINOv2, Phikon, UNI, HOptimus, and MedSigLIP, using gene-level Spearman correlation (ρ) as the primary metric. Results of predictive performance using distributional, threshold-based, and rank-based summaries are shown in Figure 1. Key points observed from these experiments are summarized in the following.

**Figure 1.**
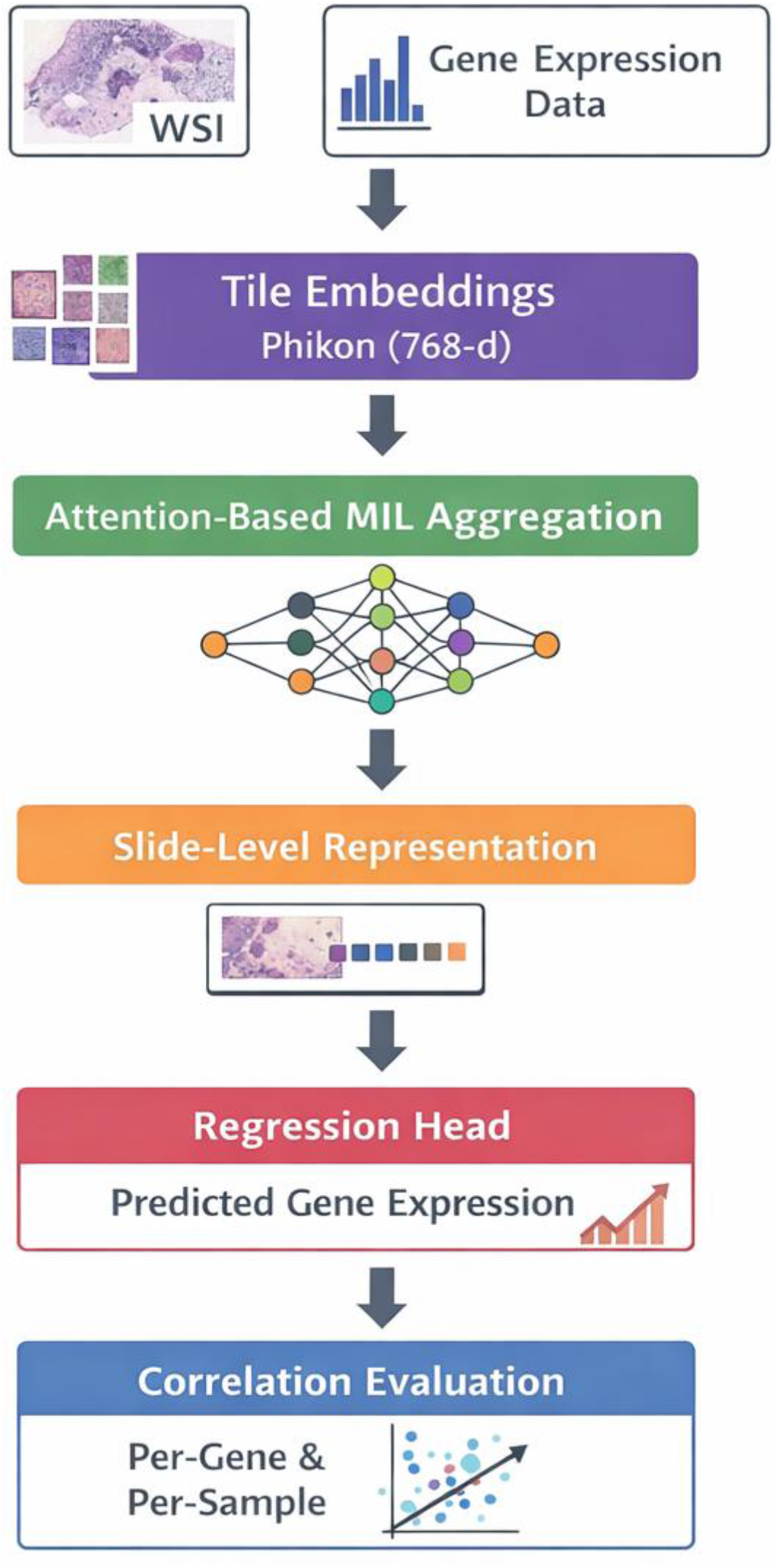
End-to-end Foundation Models Benchmarking Pipeline

**Figure 2.**
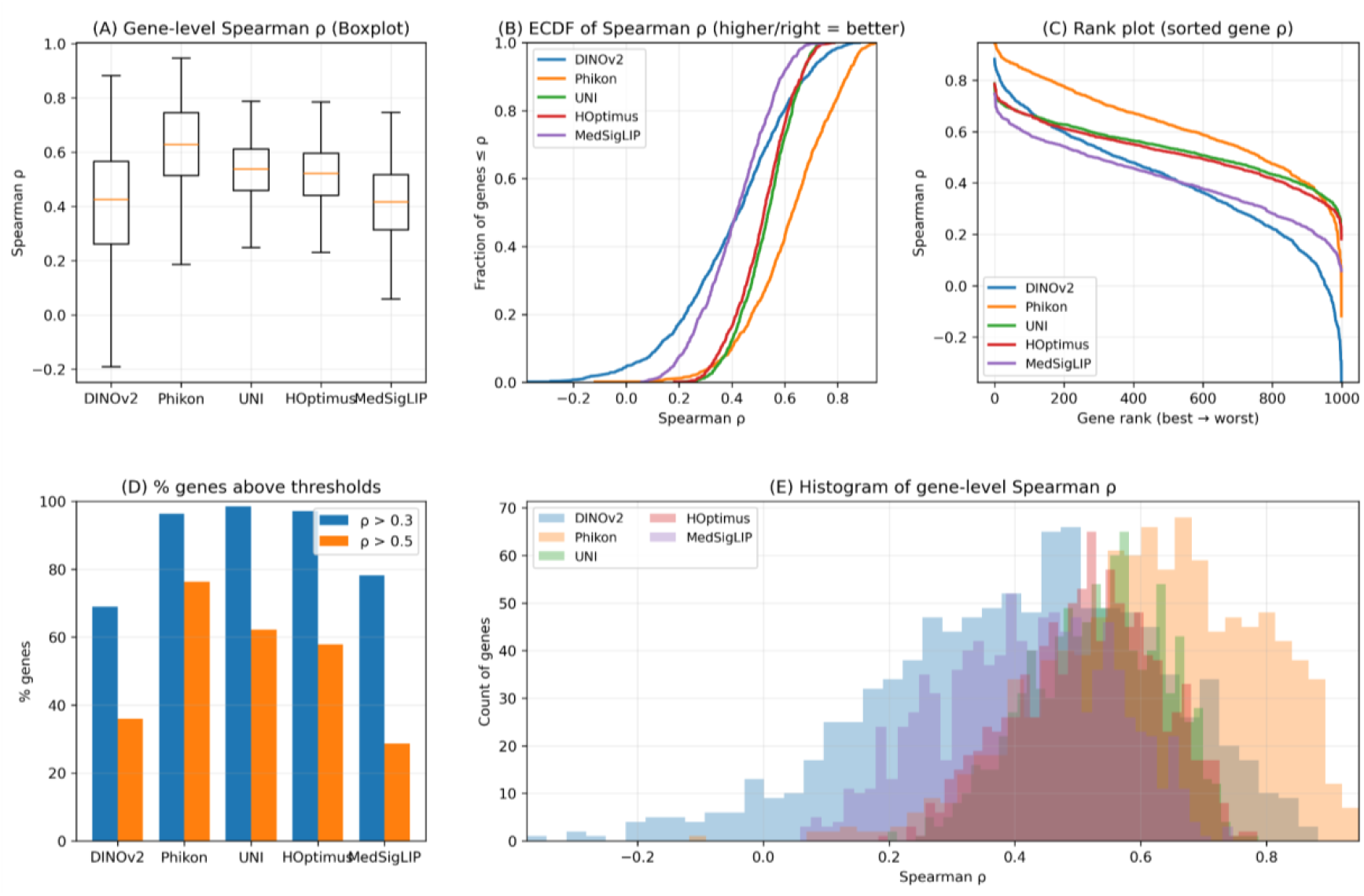
Comparative evaluation of gene expression prediction performance across five WSI feature extractors (DINOv2, Phikon, UNI, H-Optimus-0, and MedSigLIP) using gene-level Spearman correlation (ρ) as the primary metric. **(A)** Boxplot distributions of gene-level ρ across models. **(B)** Empirical cumulative distribution functions (ECDFs) of gene-level ρ highlighting the fraction of genes achieving moderate-to-high correlation. **(C)** Rank-ordered gene-level ρ curves (genes sorted by correlation) illustrating performance consistency across the full gene spectrum. **(D)** Threshold-based summary showing the percentage of genes exceeding correlation cutoffs (ρ > 0.3 and ρ > 0.5). **(E)** Histogram overlays of gene-level ρ visualizing global distributional shifts across models.

### 3.2 Gene-Level Correlation Distribution

**Figure 1(A) shows** gene-level Spearman correlation coefficient (ρ) between predicted and ground truth values of gene expression values. We observe a clear performance stratification across models. The Phikon-based model achieved the highest median correlation with a relatively compact interquartile range, indicating both strong central tendency and stability across genes. HOptimus and UNI followed with moderately high medians and comparable spread. MedSigLIP showed moderate performance, while DINOv2 exhibited the lowest median correlation and the widest dispersion, suggesting less consistent gene-level predictive capacity.

### 3.3 ECDF Analysis

**To compare the model performances beyond median values, we plotted** empirical cumulative distribution function (ECDF) curves, which can provide a global view of performance (Figure 1B). Consistent with the above observation, distribution curves shifted to the right when performance median was high and correspond to stronger overall gene-level correlations. Phikon and HOptimus display the most favorable rightward shifts, indicating a higher fraction of genes achieving moderate-to-high correlation values. Interestingly the ECDF curves were more flat for HOptimus despite their lower performance metrics being similar to Phikon and UNI, suggesting that Phikon captures expression value distributions better and performs more homogeneously for most of them, whereas the other two WSI foundation models show greater spread over performance, perhaps due to their training being biased towards some genes. DINOv2 is visibly left-shifted, confirming weaker performance across the gene spectrum.

### 3.4 Rank-Based Comparison

Rank based comparisons were made to gain more insights into model performances (Figure 1C). Ranked plots (sorted by gene-level ρ) reveal better performance consistency across the entire gene list for some models than others. Specifically, Phikon maintains superior correlation values across most ranks, particularly among best-predicted genes. HOptimus and UNI exhibit competitive mid-rank behavior but taper slightly in lower-ranked genes. DINOv2 shows a steeper decline toward lower ranks, including near-zero or negative correlations, indicating limited robustness for harder-to-predict genes.

### 3.5 Threshold-Based Evaluation

We investigated the relative proportion of genes, whose predictability exceeds predefined correlation thresholds (ρ > 0.3 and ρ > 0.5), which gives us an insight into their usability for providing a biologically interpretable outcome (Figure 1D). We observed that Phikon achieves the highest proportion of genes above both thresholds. HOptimus and UNI demonstrate strong performance at moderate thresholds (ρ > 0.3), with reduced but still competitive performance at ρ > 0.5. MedSigLIP shows moderate threshold crossing rates. DINOv2 consistently lags behind other models at both thresholds.

Overall, this analysis confirms that stronger models not only improve median correlation but also increase the number of biologically meaningful predictions.

### 3.6. Histogram of Gene-Level Correlations (Figure E)

Histogram overlays further reinforce distributional differences. Phikon’s distribution is visibly shifted toward higher correlation values, with a denser mass between 0.5–0.7. HOptimus and UNI show similar but slightly broader distributions. MedSigLIP centers around moderate correlations, while DINOv2 presents a left-skewed distribution with substantial mass below 0.3.

## 4. Discussion

The comparative performance trends indicate a consistent hierarchy: Phikon > UNI ≈ H-Optimus-0 > MedSigLIP > DINOv2 across mean ρ, median ρ, maximum ρ, and the proportion of genes exceeding ρ > 0.3 and ρ > 0.5. Phikon demonstrates superior central tendency and the highest fraction of biologically meaningful correlations, indicating stronger transcriptomic alignment at the slide level. UNI and HOptimus maintain competitive but slightly lower performance, while MedSigLIP and particularly DINOv2 underperform across all metrics.

These differences can be explained by established principles in computational pathology and representation learning. Prior studies have shown that histopathology-pretrained models outperform general vision encoders for downstream molecular tasks (Coudray et al., 2018; Kather et al., 2019; Schmauch et al., 2020). Models such as Phikon and HOptimus are pretrained directly on H&E whole-slide images, enabling them to learn tissue-specific morphological priors. This domain alignment enhances transcriptomic prediction performance.

Genomic and transcriptomic alterations manifest as systematic morphological phenotypes in H&E slides (Madabhushi & Lee, 2016; He et al., 2020). Proliferation, immune activation, stromal remodeling, and tumor microenvironmental patterns are visually encoded at cellular and architectural scales. Foundation models trained on histology are better positioned to capture these morphology-linked gene expression signatures. Although, we have not specifically carried out a functional analysis of gene sets better predicted than other, these domain-specific constructs may lie at the backbone of better prediction in histopathological images-derived foundation models compared to generic vision models. Since general-purpose models such as DINOv2 (Oquab et al., 2023) are optimized for natural image semantics and object-centric representations, they capture WSI structure information more weakly than Phikon and others. It turns out that although a large-scale self-supervised learning improves feature robustness and transferability (He et al., 2022; Dosovitskiy et al., 2021), task performance may depend not only on scale but also on domain relevance. Histopathology-specialized foundation models leverage both scale and domain-specific inductive bias, explaining their superior performance.

## 5. Conclusion

We provide a systematic benchmark of publicly available image foundation models for gene expression prediction from WSIs. Histopathology-specialized encoders consistently outperform general-purpose vision models, with Phikon achieving the strongest transcriptomic alignment in TCGA-BRCA. These findings highlight the importance of domain-aligned self-supervised pretraining for molecular inference tasks and provide practical guidance for foundation model selection in computational pathology.

